# Biomechanics of chemically treated hair for continuous 1D strands

**DOI:** 10.64898/2026.02.09.704950

**Authors:** Abhiram Podili, Allison Meer, Jash Mody, Daniel Gosnell, Alexander Vasile, Daniel Alshansky, Roche C. de Guzman

## Abstract

Human hair is a keratin-based fiber with mechanical properties relevant to load-bearing biomaterials; however, its smooth cuticle limits fiber-fiber cohesion during textile-style processing. This study examines how controlled chemical decuticularization influences surface morphology and tensile behavior of intact human hair assembled into continuous one-dimensional (1D) strands. Hair was treated with oxidative bleach, sodium hydroxide (NaOH), or formic acid (FA), carded, and spun using a standardized protocol. SEM imaging showed treatment-dependent surface disruption, from minimal cuticle modification (bleach) to partial scale lifting (NaOH) and extensive cuticle removal (FA). Tensile testing revealed significant differences in Young’s modulus, ultimate tensile strength (UTS), and elongation at break (EAB) across treatments (ANOVA, p < 0.05). NaOH-treated strands exhibited the highest modulus (207 MPa), UTS (34 MPa), and moderate extensibility (28%), whereas bleach- and FA-treated strands showed reduced stiffness and strength. Compared with reference yarns, NaOH-treated strands approached the stiffness of wool and retained greater extensibility than cotton. These findings support a processing window in which partial decuticularization enhances fiber cohesion while preserving mechanical integrity. The resulting 1D strands provide a potential building block for woven biomesh structures, motivating further evaluation of durability, cyclic behavior, multi-ply configurations, and computational modeling.

## 1. Introduction

Human hair is a keratinous fiber composed of aligned intermediate filament keratins embedded in a matrix of highly crosslinked keratin associated proteins (KAPs) (Dyer et al., 2013; Harland et al., 2022; Popescu and Hocker, 2009; Wortmann et al., 2022; Yu et al., 2017). This hierarchical structure gives rise to tensile properties comparable to certain natural and synthetic fibrous biomaterials used in load-bearing applications. The cortex, which carries most of the mechanical load, is encased by a multilayered cuticle that forms a dense, chemically resistant barrier. While this cuticle protects the cortex, its smooth, low-friction surface limits the ability of hair fibers to interlock mechanically during textile-type processes such as carding and spinning, and instead encourages “slip” between neighboring strands (Popescu and Hocker, 2009; Yu et al., 2017). As a result, intact hair is difficult to assemble into continuous, cohesive strands suitable for further downstream fabrication activities.

Many keratin-based biomaterials bypass these limitations by extracting and reconstituting keratin into hydrogels, films, or porous scaffolds (Mohamed et al., 2021; Rouse and Van Dyke, 2010). Although these processing methods allow controlled fabrication, they eliminate the native cortical microfibril architecture that contributes to the intrinsic mechanical behavior of intact hair fibers. In contrast, strategies that preserve native fiber structure while enabling assembly into yarn-like constructs remain underdeveloped. The lack of defined protocols linking chemical surface modification, fiber-scale organization, and resulting mechanical properties has limited the translation of intact hair into load-bearing or mesh-based biomaterials.

Chemical decuticularization represents a potential strategy to increase surface friction and improve spinnability. Oxidative, acidic, and alkaline treatments can lift, fragment, or partially remove cuticular layers, increasing surface roughness and promoting inter-fiber cohesion. However, aggressive treatments can also disrupt disulfide bonds, degrade keratin matrices, or damage the cortex, thereby reducing tensile performance. Previous studies show that bleaching, strong acids, and alkalis produce characteristic patterns of cuticle disruption and protein loss, yet their effects on the mechanical behavior of spun 1D hair strands remain insufficiently understood (Chen et al., 2020; Dyer et al., 2013; Franca-Stefoni et al., 2015; Velasco et al., 2022). Identifying the balance between adequate surface modification and preservation of mechanical integrity is therefore essential for developing hair-based structural biomaterials.

In this study, we examine how three chemically distinct decuticularization treatments: oxidative bleach, sodium hydroxide (NaOH), and formic acid (FA), influence both the surface morphology and the tensile properties of continuous single-ply hair strands produced through standardized carding and spinning. Surface changes were characterized using light microscopy and scanning electron microscopy (SEM). Mechanical behavior was quantified by measuring Young’s modulus, ultimate tensile strength, and elongation at break, with commercial wool, cotton, and acrylic yarns used as reference materials. Our objective is to identify a processing window in which partial cuticle removal improves fiber-fiber cohesion without substantially decreasing strand-level tensile performance. Establishing these processing-structure-property relationships is a necessary step toward the eventual fabrication of two-dimensional woven biomeshes from chemically modified human hair.

## 2. Materials and methods

### 2.1. Reagents and chemicals

Sodium dodecyl sulfate (SDS), chloroform, methanol, sodium hydroxide pellets (NaOH), and formic acid (FA, 88%-90%) were obtained from Sigma-Aldrich (St. Louis, MO, USA) and Lab Alley (Austin, TX, USA). The 40V Permanent Developer and Extra Strength Lightener powder were purchased from Clairol (Stamford, CT, USA). All reagents were used as received. Aqueous solutions were prepared with deionized water.

### 2.2. Hair collection and preparation

Human hair was obtained from local salons on Long Island, NY, via anonymized donations. Fibers were manually sorted to include strands ≥ 1 inch in length with natural dark pigmentation (eumelanin-rich). Fibers that were naturally blonde, previously bleached, colored, or visibly damaged were excluded. Non-hair debris and synthetic fibers were removed by hand. Hair strands were first washed in a 0.2% (m/V) SDS solution at warm temperature under manual agitation to remove surface contaminants. Fibers were rinsed thoroughly with water and air-dried overnight. Delipidization was then performed by immersing hair in a 2:1 V chloroform-methanol solution for 12-16 h under continuous shaking. After immersion, fibers were rinsed with water, dried with microfiber cloths, and air-dried overnight. Because prior delipidization caused excessive embrittlement when combined with NaOH, hair intended for NaOH treatment was not exposed to the chloroform-methanol step. An overview of the processing sequence from raw hair to continuous one-dimensional (1D) strands is shown in **Fig. 1**.

**Fig. 1.**
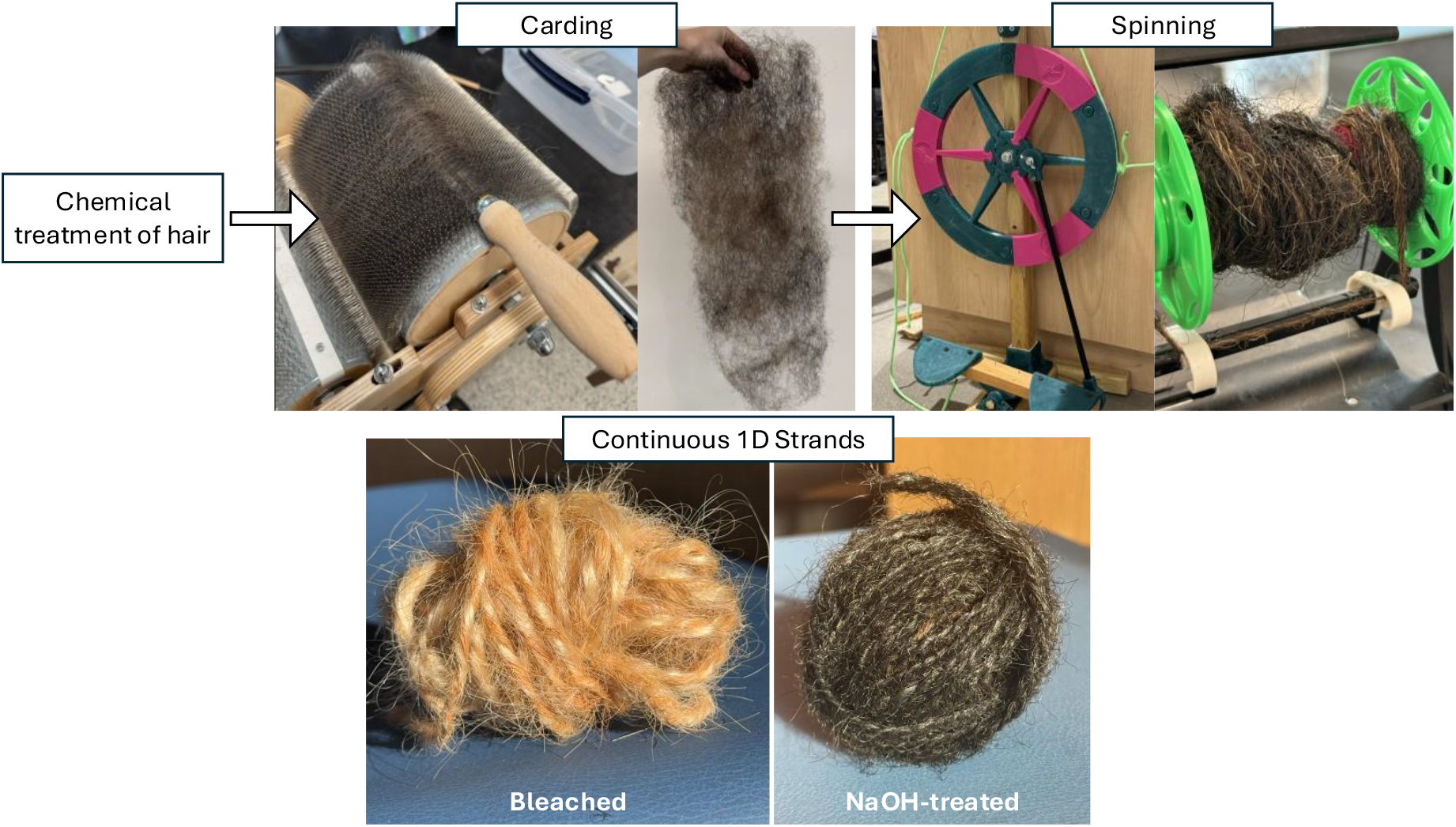
Process diagram of hair, including carding and spinning, to make the 1D strand.

### 2.3. Chemical decuticularization treatments

#### 2.3.1. Oxidative bleaching

Bleaching was conducted using a freshly prepared mixture per 1 g of hair: 2 g of 40 V developer, 1 g of lightener powder, and 35 mL deionized water. The mixture was vortexed until homogeneous and transferred into 50-mL conical tubes containing the carded hair sample. Tubes were wrapped in aluminum foil to prevent light exposure and placed on a bidirectional rotating shaker at room temperature (RT) for 24 h. Samples were washed 5× with water and air-dried. If insufficient lightening was observed, a second bleaching step was performed using half-strength developer and lightener concentrations under identical conditions.

#### 2.3.2. Formic acid treatment

Hair was processed at a ratio of 10 g of hair per 200 mL of FA solution. Hair was first boiled (at ∼100 °C) for 20 min inside a chemical fume hood, then transferred to fresh FA and incubated on a shaker at RT for 16 h. Fibers were then passed through a filter to remove small fragments, rinsed repeatedly with water, and soaked in water for an additional 24 h. Treated fibers were air-dried at ambient conditions. All formic acid handling, boiling, and waste disposal were conducted according to institutional safety protocols.

#### 2.3.3. Sodium hydroxide treatment

Untreated, non-delipidized hair samples were immersed in 1.25 M NaOH solution for 10 min at RT, then rinsed thoroughly with water and air-dried before further processing.

### 2.4. Carding and spinning into 1D strands

Following chemical treatment and drying, hair fibers were gently separated by hand to remove large tangles and brushed lightly to promote preliminary fiber alignment. Carding was performed using a drum carder (**Fig. 1**). Fiber was fed beneath the small roller in small handfuls; any material not transferred after 1-2 rotations of the drum was re-fed in subsequent passes. Once the large drum was uniformly covered, a hand carder was used to compact fibers into the bristles, allowing additional loading. When the drum was full, batts were removed with a picker tool by separating fibers at the drum gap and lifting them off in a cohesive sheet.

Carded batts were drafted manually into roving-like bundles and spun into single-ply strands using a manual spinning wheel operated under consistent twist and draw conditions (**Fig. 1**). No binders, finishes, or post-spinning chemical treatments were applied. All spun strands were conditioned at 22-24 °C and 40%-50% relative humidity for at least 24 h before testing.

### 2.5. Reference yarn materials

Commercial wool, cotton, and acrylic yarns were used as reference materials. All yarns were single-ply and of comparable nominal diameter to the spun hair strands. Reference yarns were conditioned under the same environmental conditions and were processed and tested using the same protocol as the hair strands.

### 2.6. Scanning electron microscopy

Treated and untreated hair fibers were air-dried prior to imaging. Samples were mounted on conductive stubs and sputter-coated with gold using an EMS-550 sputter coater (Electron Microscopy Sciences, Hatfield, PA, USA) to provide surface conductivity. Imaging was performed using a Q250 scanning electron microscope (FEI, Hillsboro, OR, USA) at 30 kV. Images were acquired at magnifications up to 1000× to evaluate cuticle morphology, scale disruption, and cortex exposure. Additional lower-magnification images were collected to visualize overall strand surface features where indicated.

### 2.7. Tensile testing

Single-ply 1D strands (hair and reference yarns) were cut to a gauge length of 25 cm and mounted in a uniaxial mechanical testing frame (Mark-10 Corporation, Copiague, NY, USA) equipped with grips designed to minimize slippage. A small preload was applied to remove slack prior to testing. Samples were stretched under displacement control at a constant crosshead rate of 300 cm/s until failure. Load (F)-displacement (ΔL) data were converted to engineering stress and strain using:

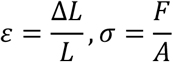

where L is the initial gauge length, and A is the initial measured cross‐sectional area. Strand diameter was measured at multiple points along the gauge length using a stereomicroscope, and the average value was used to compute A, assuming a circular cross‐section. True stress and true strain were calculated using:

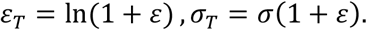

Young’s modulus (E = elastic modulus at tension) was determined from the linear elastic region of the true stress-strain curve. Ultimate tensile strength (UTS) was defined as the maximum true stress prior to failure, and elongation at break (EAB) was defined as the true strain at the point of fracture. For each material and treatment condition, at least three independent strands (n ≥ 3) were tested. Post‐failure behavior (fiber pull‐out vs. cohesive fracture) was documented qualitatively.

### 2.8. Data handling and statistical analysis

Force-displacement data were processed to obtain engineering and true stress-strain curves as described in Section 2.7. For each specimen, Young’s modulus, UTS, and EAB were extracted from the individual curves. Mechanical data are reported as mean ± 1 standard deviation (SD) for each group. Normality of the data distributions was assessed using the Shapiro-Wilk test.

One‐way analysis of variance (ANOVA) was performed to assess differences among treatment groups and between hair strands and reference yarns, with a significance level of α = 0.05. When ANOVA indicated significant differences, Tukey-Kramer post-hoc multiple comparison tests were applied. Selected pairwise comparisons were further evaluated using two-sample Student’s t-tests, with significance denoted as *p < 5%, **p < 1%, and p < 0.1%. Statistical analyses were performed in MATLAB (MathWorks, Natick, MA). Trends in mechanical behavior were interpreted in conjunction with SEM observations to relate chemical treatment, surface morphology, and strand-level performance.

## 3. Results and discussion

### 3.1. Macroscopic characteristics of continuous 1D strands

All chemically treated and untreated hair samples could be carded and spun into continuous single-ply 1D strands using the standardized workflow (**Fig. 1**). Representative macroscopic and stereomicroscope images (**Fig. 2**) show qualitative differences in cohesion and uniformity among treatment groups. NaOH-treated strands exhibited comparatively uniform diameter along their length with limited fraying, consistent with enhanced fiber-fiber cohesion during spinning. Untreated strands appeared continuous but showed more local variations in thickness and loose surface fibers. FA-treated strands showed visible fragmentation and surface irregularities, with localized thinning and broken fiber ends, reflecting the more aggressive cuticle and cortex damage seen in microscopy. Bleach-treated strands were intermediate in appearance, retaining continuity but with less apparent cohesion than NaOH-treated strands. These observations indicate that all treatment conditions can be processed into 1D strands, but the resulting macroscopic quality depends on the balance between surface modification and fiber integrity.

**Fig. 2.**
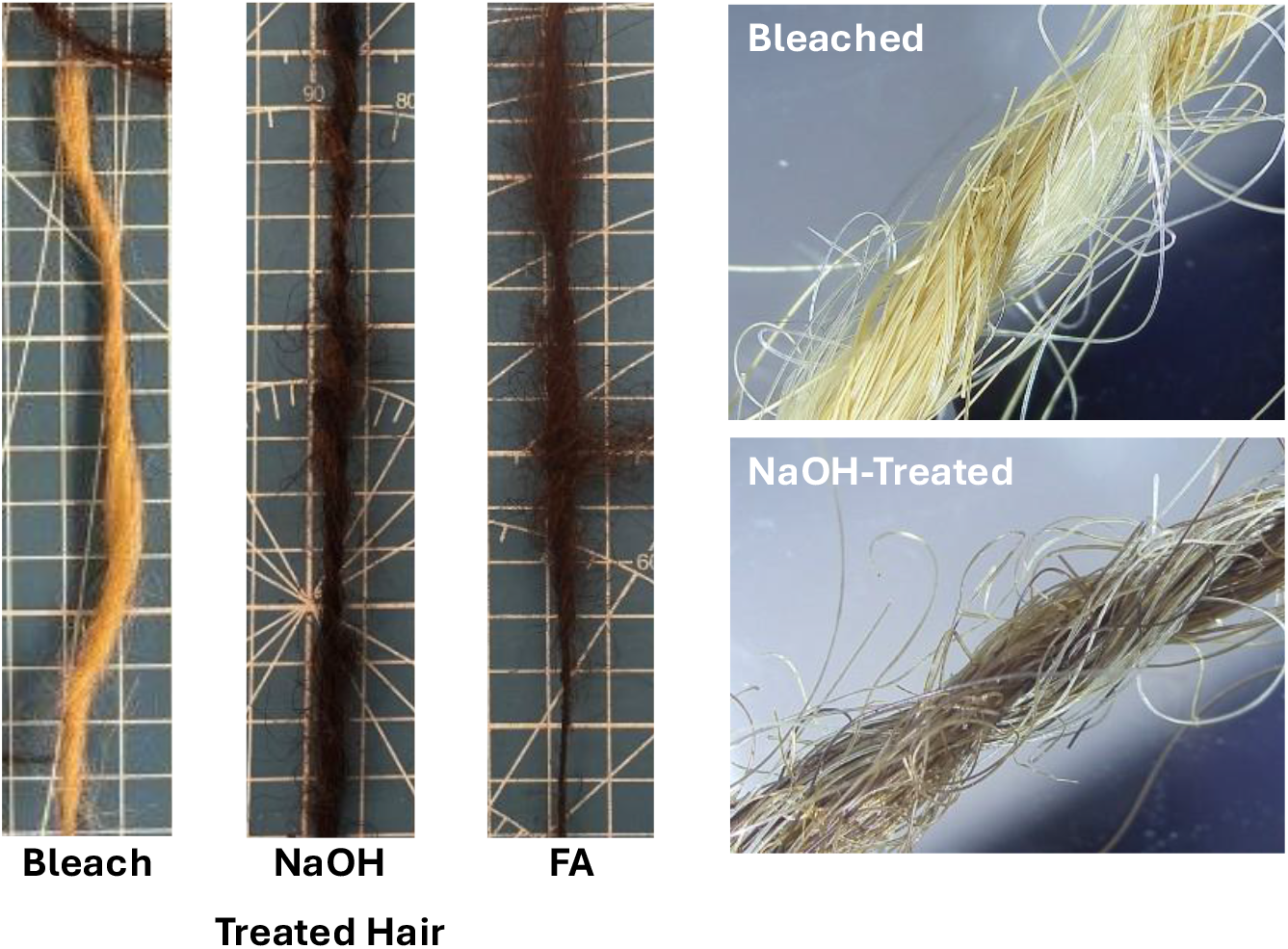
Representative samples of carded and spun hair into 1D strands.

### 3.2. Microscopic analysis of individual hair fibers and cuticle structure

#### 3.2.1. Light microscopy

Light transmission and stereomicroscopy of individual fibers (**Fig. 3**) revealed treatment-dependent changes in cuticle integrity and apparent cortex continuity. FA-treated fibers showed extensive disruption of the outer surface, with irregular contours and regions lacking discernible cuticle structure, consistent with substantial chemical degradation. In contrast, fibers from NaOH-treated and untreated groups maintained more regular cylindrical geometry, with NaOH-treated fibers exhibiting roughened surfaces consistent with partial cuticle lifting rather than complete removal.

**Fig. 3.**
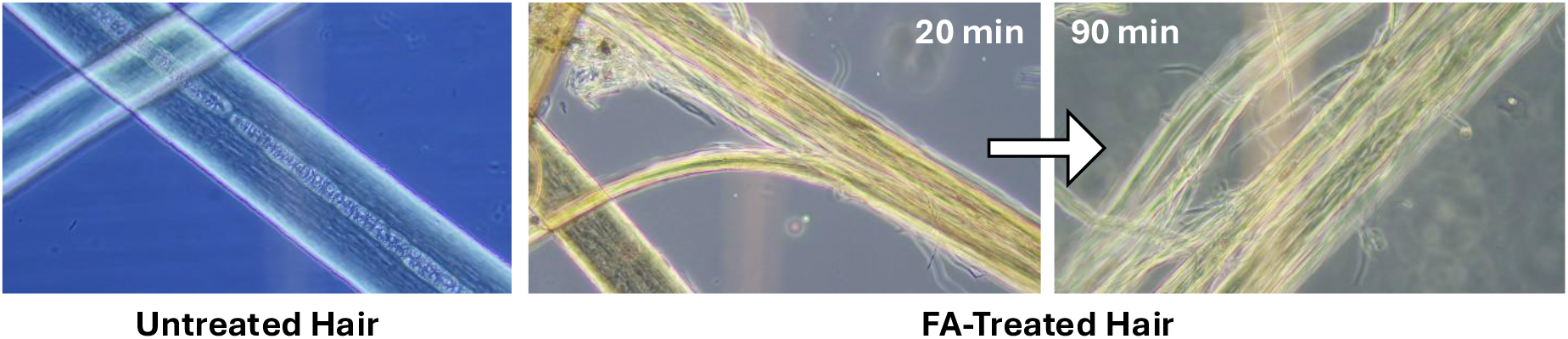
Analysis of individual fibers via light microscopy, showing the formic acid (FA) treatments.

#### 3.2.2. Ultrastructure using electron microscopy

SEM imaging of untreated human hair, sheep wool, and cat hair (**Fig. 4**) provided a broader context for cuticle structure across keratin-based fibers. All three exhibited overlapping scale-like cuticle cells; however, human hair showed relatively smoother, more closely apposed scales than wool, which displayed more pronounced scale edges, consistent with its well-known felting behavior. This comparison supports the premise that enhancing surface roughness in human hair could improve fiber-fiber interlocking in textile-like applications.

**Fig. 4.**
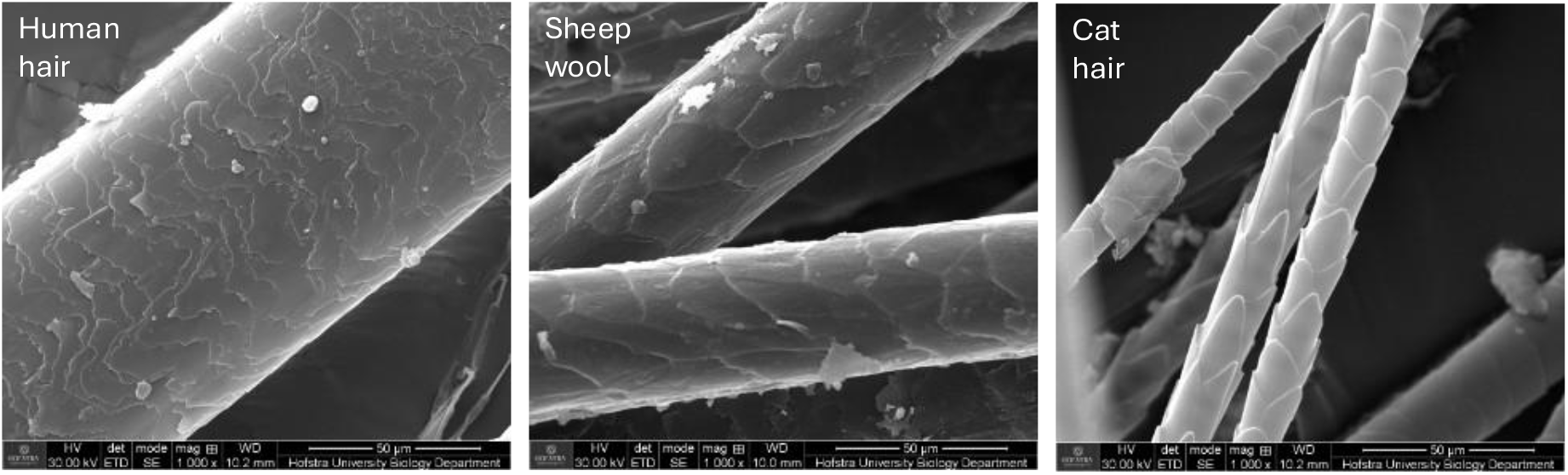
Electromicrographs of untreated human hair compared to sheep wool and cat hair.

SEM of chemically treated human hair (**Fig. 5**) revealed a continuum of cuticle disruption. Untreated fibers maintained intact, overlapping scales with smooth surfaces, indicating that SDS washing and organic solvent delipidization alone did not significantly alter cuticular structure. Bleach-treated fibers showed mild cuticle “flaking,” with some lifting and rounding of scale edges but largely preserved scale architecture. NaOH-treated fibers exhibited clear partial lifting, separation, and occasional detachment of cuticle cells, producing a rougher surface and increased topographical contrast. FA-treated fibers demonstrated the most severe damage, with near-complete loss of recognizable scale morphology and direct exposure of the cortex, along with surface pitting and fragmentation.

**Fig. 5.**
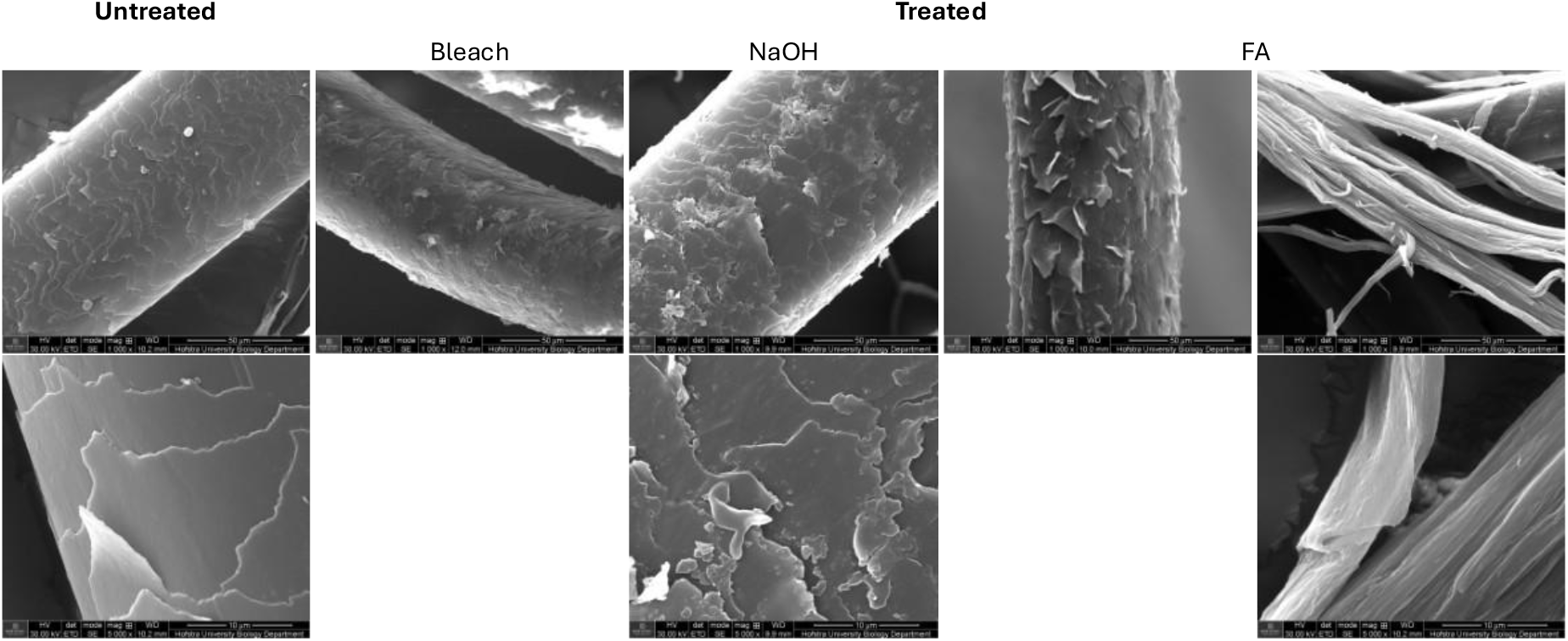
Representative SEM images of untreated versus chemically-treated human hair: bleach, NaOH, and FA.

Taken together, light microscopy and SEM show that the three chemical treatments span a range from minimal surface modification (bleach) to partial decuticularization with preserved cortex (NaOH) and extensive cuticle removal with likely cortical damage (FA). This morphological spectrum provides a structural framework for interpreting the differences in tensile behavior among groups.

### 3.3. Tensile behavior of spun hair strands

Representative true stress-strain curves for untreated and chemically treated hair strands, alongside commercial acrylic, cotton, and wool yarns, are shown in **Fig. 6**. All groups exhibited an initial linear region followed by nonlinear deformation and failure; however, the breadth and shape of the curves differed substantially among materials. These differences reflect variation in fiber packing density, degree of structural uniformity, and failure mode.

**Fig. 6.**
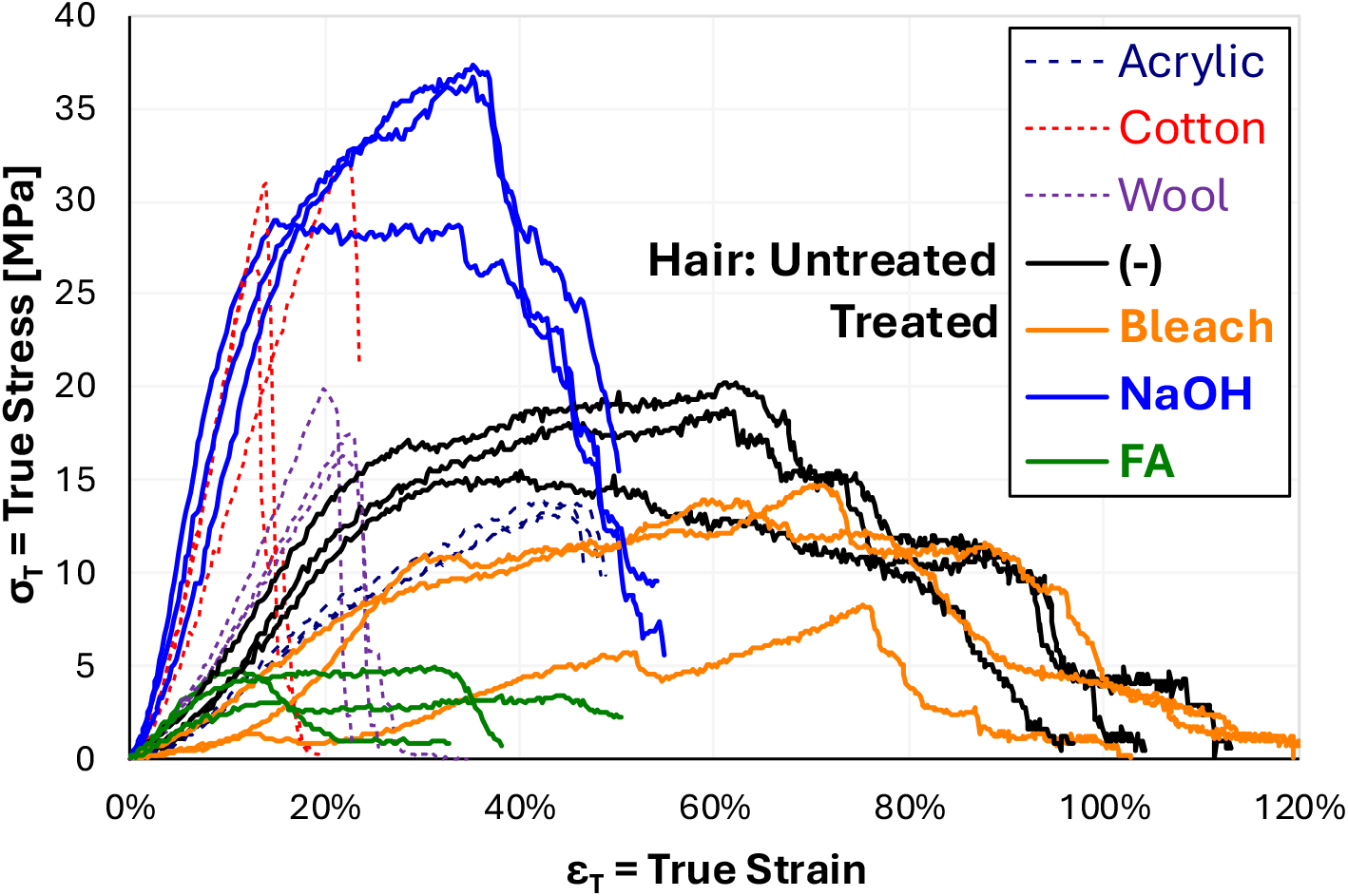
Individual tensile stress-strain data of untreated versus treated human hair as 1D strands in comparison with reference yarn materials: wool, cotton, and acrylic.

Commercial acrylic, cotton, and wool yarns showed narrow, highly consistent stress-strain profiles with low variability (**Table 1**). Their tight packing of many fine, uniformly processed fibers produces a highly integrated structure, allowing individual filaments to share load efficiently. During failure, the individual fibers in these yarns fracture nearly simultaneously, leading to a steep rise to peak stress followed by a sharp drop in the stress-strain curve. This simultaneous failure explains their low coefficient of variation (3%-18%) and narrow elongation ranges.

**Table 1.** Variability within samples: 1 standard deviation/mean.

| Groups | E | UTS | EAB |
| --- | --- | --- | --- |
| Acrylic | 5% | 3% | 5% |
| Cotton | 24% | 10% | 31% |
| Wool | 18% | 7% | 10% |
| (-) | 16% | 14% | 92% |
| Bleach | 54% | 29% | 10% |
| NaOH | 24% | 13% | 41% |
| FA | 32% | 19% | 58% |

In contrast, the lab-made 1D hair strands exhibit broader and more variable stress-strain curves. Unlike commercial yarns, spun hair strands contain fewer fibers, larger diameters, and less uniform packing. As a result, individual fibers do not fail simultaneously but rather break progressively along the gauge length. This staggered failure produces an extended nonlinear region and broad strain ranges, particularly for untreated and bleach-treated strands. Untreated and bleached hair showed the highest variability, with failure strains spanning approximately 30%-120%. Their smooth, intact cuticles promote fiber-fiber slippage, reducing synchronized load sharing and causing individual fibers to fail independently at different points in the test. Bleach additionally introduces oxidative defects that increase heterogeneity along the fiber length, further broadening strain variability. Formic acid-treated strands exhibited intermediate variability, with strains ranging roughly 10%-50%. Although FA removes most of the cuticle and increases surface roughness, the associated cortical degradation and fragmentation create weak points along the fiber, limiting extensibility but still producing asynchronous fiber failures. NaOH-treated strands demonstrated the narrowest strain range among the chemically treated groups (20%-50%) and the most cohesive failure behavior. Partial cuticle lifting increases friction and reduces inter-fiber slippage, while the cortical structure remains largely intact. This combination promotes more uniform load distribution across fibers, resulting in more synchronized failure events and reduced variability compared with other hair groups.

Across all mechanical parameters (**Fig. 7**): Young’s modulus, ultimate tensile strength (UTS), and elongation at break (EAB), NaOH-treated strands showed the strongest and most consistent performance, with statistically significant differences from bleach and FA groups (ANOVA with Tukey-Kramer, p < 0.05; selected pairwise t-tests shown in **Table 2**). Their modulus approached that of wool yarn, while maintaining greater extensibility than cotton, placing them within the mechanical range of keratin-based textile fibers.

**Table 2.** Student’s t-test p-values of pairwise comparison among groups and significance at * < 5%, ** < 1%, and *** < 0.1%.

| Pairs |  | E | UTS | EAB | E | UTS | EAB |
| --- | --- | --- | --- | --- | --- | --- | --- |
| Acrylic | Cotton | 0.50% | 0.06% | 0.10% | ** | *** | *** |
|  | Wool | 1.32% | 0.60% | 0.04% | * | ** | *** |
|  | (-) | 3.45% | 3.42% | 59.01% | * | * |  |
|  | Bleach | 21.89% | 55.89% | 0.34% |  |  | ** |
|  | NaOH | 0.46% | 0.15% | 8.07% | ** | ** |  |
|  | FA | 93.47% | 0.01% | 18.91% |  | *** |  |
| Cotton | Wool | 1.29% | 0.22% | 8.66% | * | ** |  |
|  | (-) | 0.85% | 0.53% | 39.03% | ** | ** |  |
|  | Bleach | 0.45% | 0.24% | 0.04% | ** | ** | *** |
|  | NaOH | 62.90% | 25.70% | 17.26% |  |  |  |
|  | FA | 0.56% | 0.01% | 28.30% | ** | *** |  |
| Wool | (-) | 21.43% | 71.50% | 60.36% |  |  |  |
|  | Bleach | 1.77% | 7.27% | 0.04% | * |  | *** |
|  | NaOH | 1.03% | 0.36% | 52.42% | * | ** |  |
|  | FA | 3.72% | 0.01% | 62.45% | * | *** |  |
| (-) | Bleach | 4.10% | 7.84% | 12.34% | * |  |  |
|  | NaOH | 0.72% | 0.58% | 79.37% | ** | ** |  |
|  | FA | 11.68% | 0.08% | 82.05% |  | *** |  |
| Bleach | NaOH | 0.41% | 0.28% | 0.60% | ** | ** | ** |
|  | FA | 34.88% | 1.95% | 1.76% |  | * | * |
| NaOH | FA | 0.50% | 0.04% | 97.37% | ** | *** |  |

**Fig. 7.**
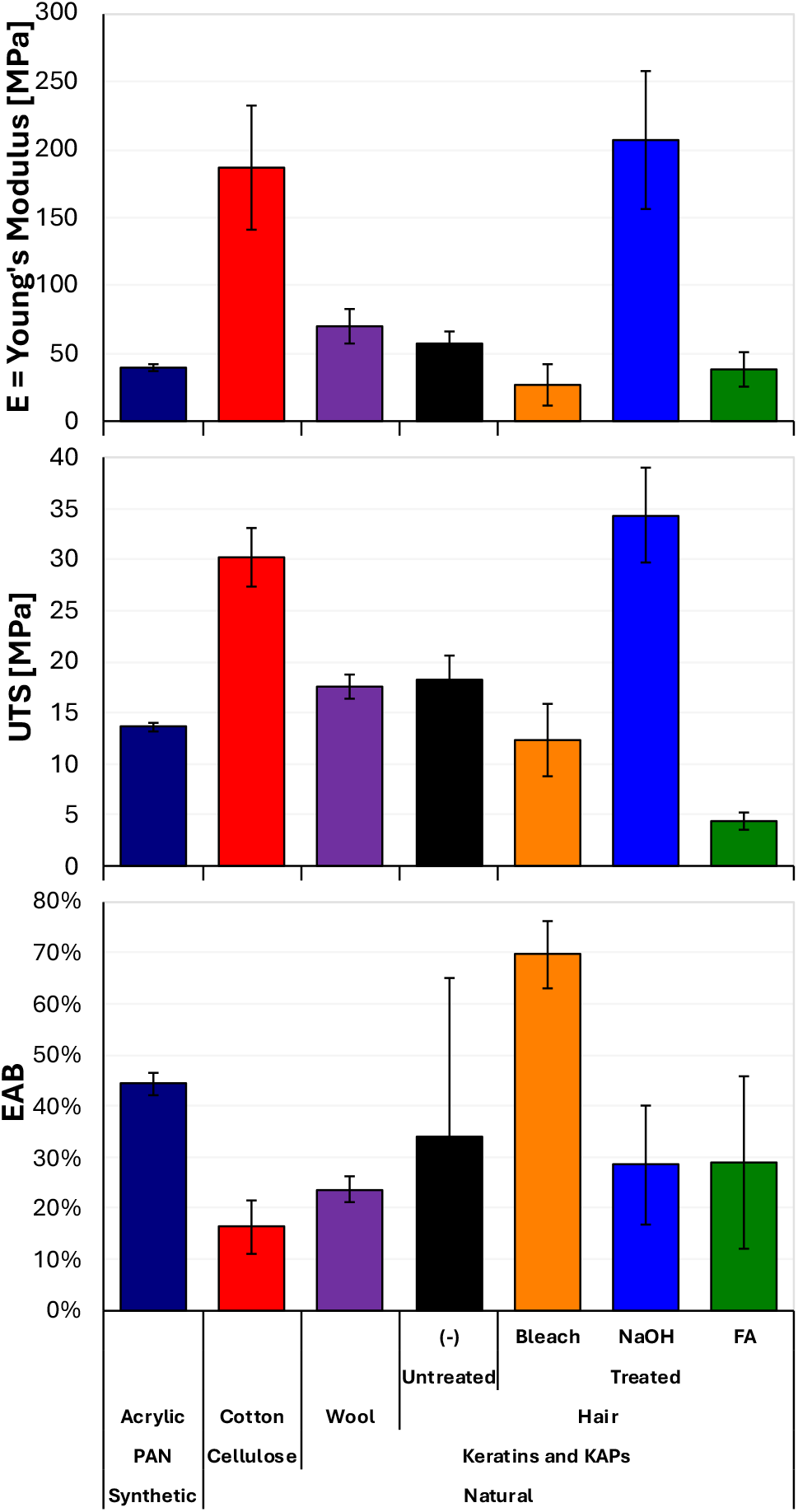
Biomechanical properties: Young’s modulus (E), ultimate tensile strength (UTS), and elongation at break (EAB) of untreated versus chemically treated 1D hair strands which are natural biomaterials made from keratins and KAPs. In comparison, reference yarn materials: synthetic polyacrylonitrile (PAN) acrylic, natural: cotton (cellulose) and sheep wool (keratins and KAPs).

Overall, the mechanical behavior of the 1D strands is governed by the interplay between chemical modification, surface friction, fiber packing density, and failure synchrony. Commercial yarns fail coherently due to tightly packed, uniform filaments. Lab-made hair strands, particularly untreated and bleached samples, fail progressively due to looser packing and heterogeneous microstructure. NaOH treatment uniquely shifts the behavior toward more coherent load sharing without sacrificing extensibility, indicating its suitability for textile-style fabrication and potential incorporation into 2D woven biomeshes.

### 3.4. Linking surface modification, cohesion, and mechanical performance

The relationship between chemical surface modification and the mechanical behavior of the resulting 1D strands reflects the balance between increased fiber-fiber friction and preservation of load-bearing capacity. Partial cuticle lifting achieved through NaOH treatment provided the most effective combination of these features. The increased surface roughness improved cohesion during drafting and spinning, while the underlying cortex remained structurally intact, supporting higher modulus and strength. In addition, NaOH-treated strands showed narrower strain ranges (approximately 20%-50%) compared with untreated, bleach, and FA groups, indicating more synchronized fiber failure and improved load sharing.

Untreated and bleached hair retained smoother cuticles and therefore lower friction between fibers. During tensile loading, individual fibers slipped and failed progressively rather than collectively, resulting in broad strain spreads (30%-120% for untreated and bleached) and lower reproducibility. In contrast, FA treatment produced extensive cuticle removal and cortex damage, increasing surface friction but reducing mechanical integrity. FA-treated strands exhibited fragmented surfaces and strain ranges of ∼10%-50%, reflecting premature failure initiated at chemically weakened regions.

Compared with reference yarns, commercial acrylic, cotton, and wool exhibited tightly clustered stress-strain curves and minimal variability due to densely packed, uniformly aligned filaments that fail nearly simultaneously. The broader spread in lab-made hair strands reflects their lower packing density and heterogeneous fiber properties. Among the chemical treatments, NaOH-treated strands most closely approached the mechanical behavior of wool, demonstrating that controlled partial decuticularization improves both cohesion during spinning and mechanical consistency during tensile loading.

### 3.5. Limitations and future work

Several limitations should be considered. The donor hair used in this study varied in diameter, age, and prior environmental exposure, all of which may contribute to scatter in the mechanical data. Although chemical treatment reduced some variability, the natural heterogeneity of human hair remains a challenge for consistency in strand fabrication. Mechanical testing was limited to uniaxial tension of single-ply strands, which does not capture fatigue behavior, creep, stress relaxation, torsional resistance, or abrasion performance, properties that are relevant to textile-style processing and woven mesh applications.

Future work should evaluate multi-ply strand configurations, where additional plies may increase packing density and reduce asynchronous fiber failure. Quantifying bending stiffness, cyclic response, and abrasion resistance will provide a more complete understanding of durability under expected handling and use conditions. Finite element analysis (FEA) could help model stress distributions along strands, particularly in regions of partial cuticle disruption, and may guide optimization of chemical treatment parameters. Integration of optimized strands into 2D woven structures will be essential for assessing load transfer across intersections, scalability, and suitability for biomedical or structural applications.

## 4. Conclusions

This study demonstrates that chemical surface modification of intact human hair influences both fiber-fiber cohesion during spinning and the mechanical behavior of resulting 1D strands. Among the treatments tested, NaOH produced controlled partial cuticle lifting while preserving cortical integrity, resulting in increased stiffness, greater strength, and reduced variability compared with untreated, bleached, and FA-treated hair. The tensile behavior of NaOH-treated strands approached that of wool while maintaining higher extensibility than cotton, indicating that human hair can be tuned to fall within the mechanical range of established keratin-based yarns.

These findings support a processing window where moderate decuticularization enhances spinnability and inter-fiber friction without compromising mechanical performance. Although further work is needed to evaluate multi-ply structures, durability under cyclic loading, and woven 2D architectures, the results establish a foundation for developing hair-based biomaterials with mechanical properties suitable for textile-like and potentially biomedical applications.

## CRediT authorship contribution statement

**Abhiram Podili:** Investigation, Formal analysis, Writing – original draft. **Allison Meer:** Conceptualization, Methodology, Investigation. **Alexander Vasile:** Investigation, Formal analysis, Writing – original draft. **Jash Mody:** Investigation, Visualization, Writing – original draft. **Daniel Gosnell:** Investigation, Writing – original draft. **Daniel Alshansky:** Investigation, Writing – original draft. **Roche C. de Guzman:** Conceptualization, Methodology, Resources, Formal analysis, Visualization, Data curation, Writing – original draft, Writing – review & editing.

## Acknowledgement

We thank the members of Dr. de Guzman’s Bioengineering Materials Lab and the Biology Department, as well as Jason Williamns for his help with the SEM. We also appreciate Fusion Beauty Salon, Stylush Salon and Laser, Fresh Cuts, and Hair Designers for providing human hair samples.

